## Supplementary material for "Midbody proteins display distinct temporal dynamics during cytokinesis": This PDF file includes: Figure S1, S2 and associate legends; Legend for Supplementary Table S1; Legends for Supplementary Videos S1 to S4

- Figure S1, S2 and associate legends
- Legend for Supplementary Table S1
- Legends for Supplementary Videos S1 to S4

**Other Supplementary Information for this manuscript include the following:**

- Videos S1 to S4
- Supplementary Table S1

**Supplementary Figures**

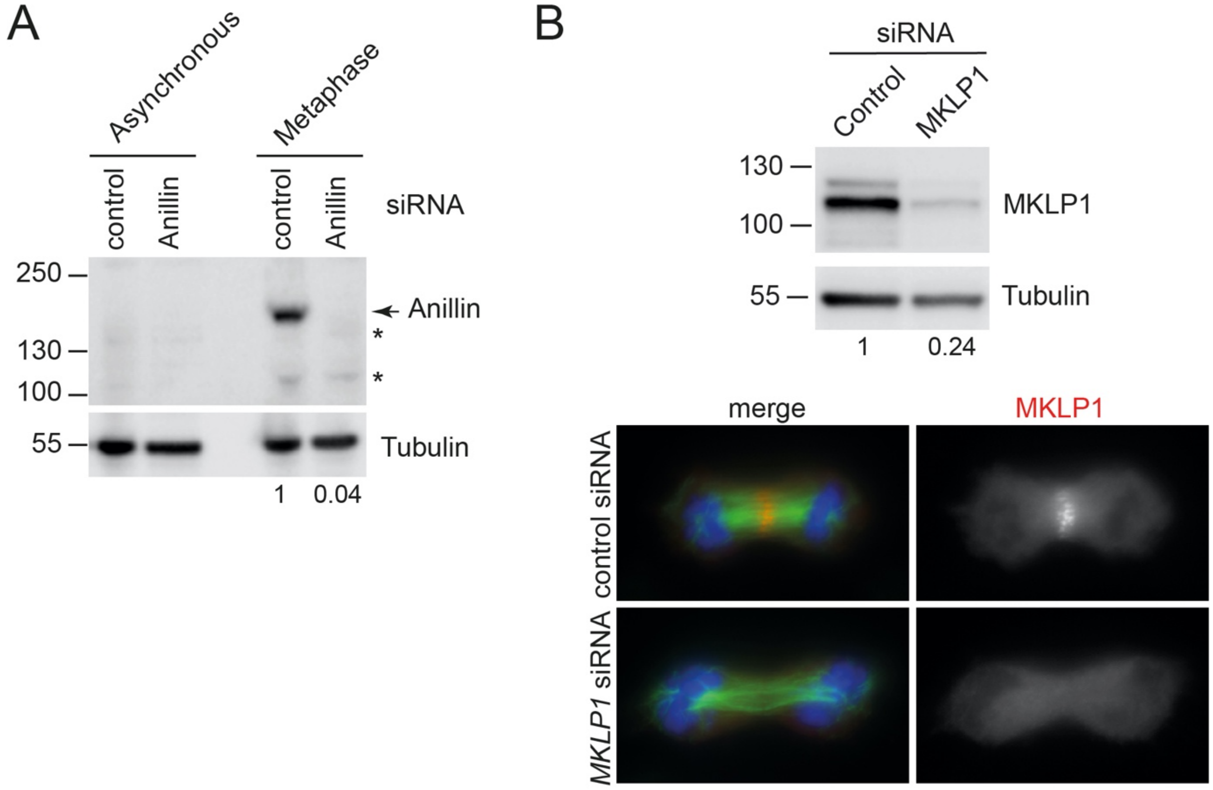


**Figure S1.** Validation of Anillin and KIF23/MKLP1 antibodies. (A) HeLa Kyoto cells were treated with siRNAs directed against either a random sequence (control) or Anillin (ANLN). After 48 hours asynchronous cells or cells synchronized in metaphase by thymidine/nocodazole block were collected and proteins were extracted and analyzed by western blot to detect Anillin and tubulin (loading control). The numbers on the left indicate the sizes in kDa of the molecular mass marker. The number at the bottom indicate the level of Anillin normalized to tubulin and relative to the control. The asterisks marks non-specific bands. (B) Top, HeLa Kyoto cells were treated with siRNAs directed against either a random sequence (control) or KIF23/MKLP1. After 48 hours cells were collected and proteins were extracted and analyzed by western blot to detect KIF23/MKLP1 and tubulin (loading control). The numbers on the left indicate the sizes in kDa of the molecular mass marker. The number at the bottom indicate the level of KIF23/MKLP1 normalized to tubulin and relative to the control. Bottom, HeLa cells were treated with siRNAs directed against either a random sequence (control) or KIF23/MKLP1 and after 48 hours were fixed and stained to detect DNA (blue), tubulin (green) and KIF23/MKLP1 (red). Note that *KIF23/MKLP1* siRNA cells show a feeble and disorganized central spindle. Bars, 10 μm.

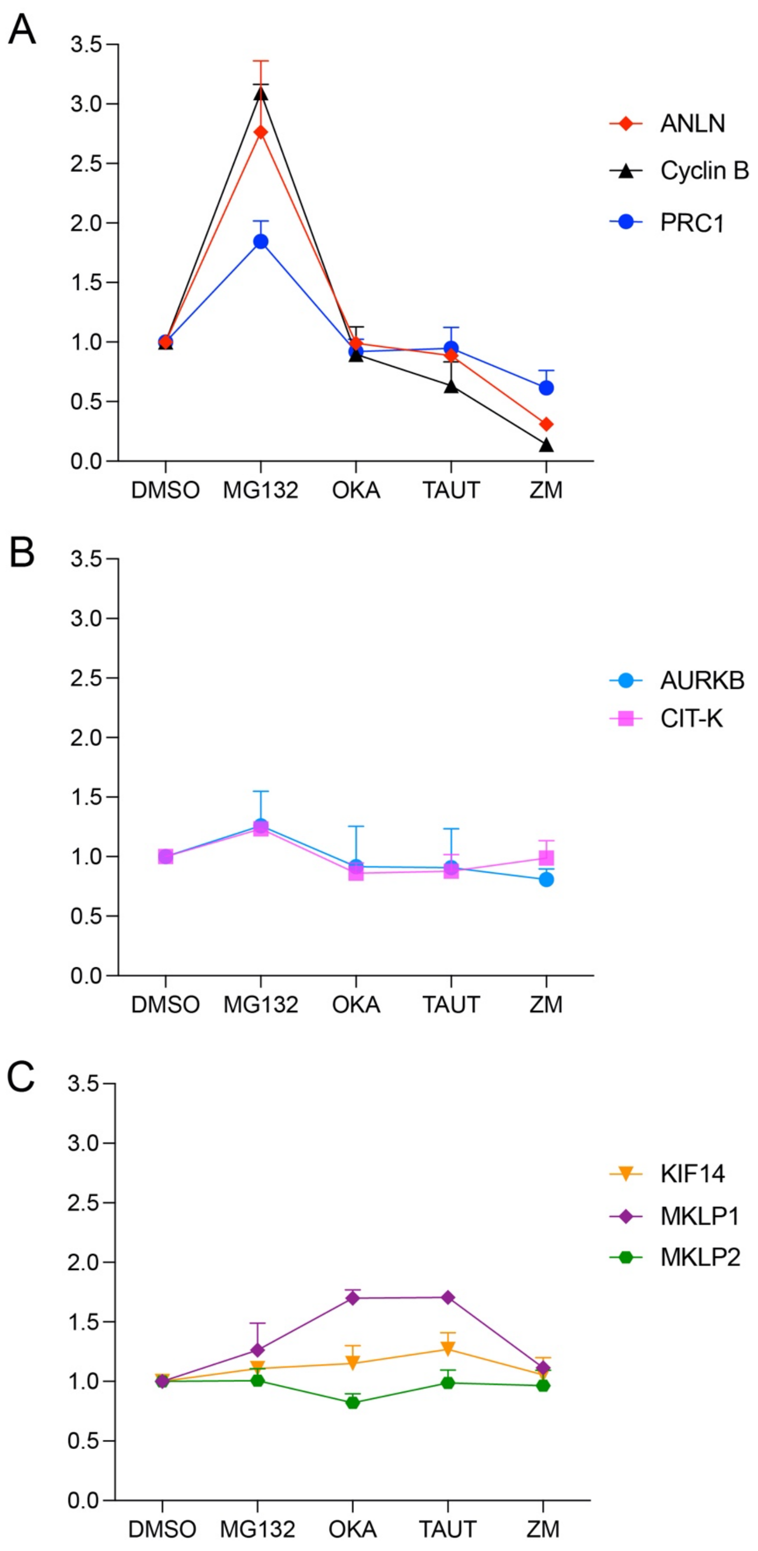


**Figure S2.** The levels of midbody proteins are differentially affected by treatments with inhibitors of the proteasome and mitotic kinases and phosphatases. The data shown in the graph in Fig. 2D were separated in three groups based on their response to drug treatments.

### Supplementary Video Captions

**Supplementary Video S1.** Time-lapse analysis of AURKB dynamics during cytokinesis. This movie shows the dynamics of GFP::AURKB and chromosomes, visualized using SiR-DNA, during cytokinesis; see Fig. 3A for details. All sequences were captured at 2 min intervals. Playback rate is 5 frames per second (FPS).

**Supplementary Video S2.** Time-lapse analysis of PRC1 dynamics during cytokinesis. This movie shows the dynamics of GFP::PRC1 and chromosomes, visualized using SiR-DNA, during cytokinesis; see Fig. 3B for details. All sequences were captured at 2 min intervals. Playback rate is 5 frames per second (FPS).

**Supplementary Video S3.** Time-lapse analysis of CIT-K dynamics during cytokinesis. This movie shows the dynamics of CIT-K::GFP and chromosomes, visualized using SiR-DNA, during cytokinesis; see Fig. 3C for details. All sequences were captured at 2 min intervals. Playback rate is 5 frames per second (FPS).

**Supplementary Video S4.** Time-lapse analysis of KIF23/MKLP1 dynamics during cytokinesis. This movie shows the dynamics of MKLP1::GFP and chromosomes, visualized using SiR-DNA, during cytokinesis; see Fig. 3D for details. All sequences were captured at 2 min intervals. Playback rate is 5 frames per second (FPS).

**Supplementary Table Caption**

**Table S1.** List of the proteins present in the midbody ubiquitylation interactome. Excel file listing the proteins identified in the midbody ubiquitylation interactome along with their respective gene names and GO terms. Baits are in red. The proteins found to be specifically associated with either ‘transient’ or ‘stable’ midbody proteins are shown in separate worksheets. Note that the ‘transient’ and ‘stable’ midbody ubiquitylation interactomes were generated using a reduced number of midbody bait proteins compared to the entire midbody ubiquitylation network shown in the first sheet and in Figure 4A.
